# Acyclovir Improves the Efficacy of Chemoradiation in Nasopharyngeal Cancer Containing the Epstein Barr Virus Genome

**DOI:** 10.1101/2020.10.18.343236

**Authors:** Aditya Thandoni, Andrew Zloza, Devora Schiff, Malay Rao, Kwok-wai Lo, Bruce G. Haffty, Sung Kim, Sachin R. Jhawar

## Abstract

Nasopharyngeal carcinoma (NPC) is a malignancy endemic to East Asia and is caused by Epstein-Barr Virus (EBV)-mediated cancerous transformation of epithelial cells. The standard of care treatment for NPC involves radiation and chemotherapy. While treatment outcomes continue to improve, up to 50% of patients can be expected to recur by five years, and additional innovative treatment options are needed. We posit that a potential way to do this is by targeting the underlying cause of malignant transformation, namely EBV. One method by which EBV escapes immune surveillance is by undergoing latent phase replication, during which EBV expression of immunogenic proteins is reduced. However, chemoradiation is known to drive conversion of EBV from a latent to a lytic phase. This creates an opportunity for the targeting of EBV-infected cells utilizing anti-viral drugs. Indeed, we found that combining acyclovir with cisplatin and radiation significantly decreases the viability of the EBV-infected C666-1 cell line. Western blot quantification revealed a resultant increase of thymidine kinase (TK) and apoptosis-inducing mediators, cleaved PARP (cPARP) and phosphorylated ERK (pERK). These studies suggest that the addition of anti-viral drugs to frontline chemoradiation may improve outcomes in patients treated for EBV-related NPC and future *in vivo* and clinical studies are needed.

## Introduction

Nasopharyngeal carcinoma (NPC) is a rare malignancy in most of the world, but is endemic in southern China and parts of Southeast Asia^1, 2^. Almost all cases of endemic NPC are associated with mucosal epithelial cell infections with the Epstein-Barr Virus (EBV)^1^, thus establishing a crucial association between the virus and the induction of transformed and invasive cancer^2^. EBV is a latency type II gamma herpes virus containing a genome of nearly 100 genes, with a majority of gene expression present during the lytic phase, compared to only eleven viral genes present during the latent phase^3^. Although naturally immunogenic, the ability of EBV to transition to latent phase replication provides a means of escape from immune surveillance through the decrease in expression of immunogenic proteins via epigenetic mechanisms^4^. While nucleoside-analogue anti-herpesvirus drugs, like acyclovir, can work in the lytic phase, leading to the death of infected cells (including tumor cells), these drugs remain in their inactive prodrug form in the latent phase, making them ineffective as primary therapy for EBV-associated malignancies^5, 6^. The role of EBV in NPC is important, as plasma titers of EBV DNA in NPC patients pre and post treatment correlate with the stage of NPC, tumor burden in patients, and changes in titers can predict outcomes^7^.

The backbone of treatment of advanced NPC is the utilization of chemotherapy and radiation. Studies have shown that each of these two therapies may individually function as a lytic phase inducer of EBV. Therefore, we sought to determine whether targeting EBV in the context of chemoradiation may augment *in vitro* signatures of treatment response, including decreased tumor cell viability and increased anti-tumor pathway signaling.

## Methods

### Tumor cell lines

C666-1, an EBV-infected nasopharyngeal cancer (NPC) human cell line, and HK-1, a non-EBV infected human NPC cell line were utilized in cell viability and western blot studies. Dr. Kwok-wai Lo and The Chinese University of Hong Kong generously donated both cell lines to The Cancer Institute of New Jersey.

### Cell viability assay

The non-radioactive colorimetric tetrazolium (MTT) assay, measuring cellular metabolic activity, was utilized as an indicator of cell viability, as per Sigma-Aldrich instructions. C666-1 cells were placed at 10,000 cells per each well of a 96-well flat-bottom plate. Experimental arms were repeated in 8 wells per treatment. The cells were cultured for 12-15 hours in Gibco’s RPMI media supplemented with 1x Glutamax, 10% FBS, and 1% penicillin -steptomycin. Cells were left untreated (no treatment; NT) or treated with acyclovir (200ug/ml), cisplatin (cis; 1ug/ml), and radiation (RT; 0-8 Gy) alone and in combinations, as described within the results, figures, and figure legends for each experiment shown. Cells were dosed with their respective drug concentrations 24 hours after initially plating. RT was conducted 24 hours after the administration of drug treatment using the Gamma Cell 40 Exactor (MDS Nordion) irradiator. Colorimetric changes in the media were detected using the Tecan Infintie M200 Pro plate reader at a wavelength of 570nm.

### Western blot analysis

Western blot analysis, as described by the protocol set by Invitrogen, was used to determine the expression of proteins of interest, including, cleaved PARP (cPARP; 1uL:2ml in PBS), phosphorylated AKT (pAKT; 1uL:2ml in PBS) phosphorylated ERK (pERK; 1uL:2ml in PBS), and thymidine kinase (TK; 1uL:2ml in PBS). GAPDH was utilized as a loading control. Quantification was conducted utilizing ImageJ software (version 1.53a; NIH), using the GELS analysis feature, as previously described^8^. All primary antibodies were obtained from CellSignaling. Secondary anti-rabbit antibody (1uL:5mL in PBS) was also obtained from CellSignaling.

### Statistical Analysis

To determine statistical significance for data comparisons, 1-way ANOVA with Tukey correction (for comparing the means of each group to every other group within an experiment) or Dunnet correction (for focused comparisons of one group to all other groups). Prism version 8.0 (GraphPad) was used for generation of all graphs and performance of statistical analyses. Bar graphs display the mean, and error bars represent the standard error of the mean (S.E.M.). Statistical significance is denoted as ns, not significant, \**P* < 0.05, \*\**P* < 0.01, \*\*\**P* < 0.001, and \*\*\*\**P* < 0.0001.

## Results

The cell viability assays demonstrate a significant increase in cell death of the C666-1 line in the presence of both acyclovir and cis compared to cells that received no treatment, acyclovir alone, and cis alone (Figure 1). Cell viability in the combination group decreased significantly as radiation fraction size increased. Western blot analysis established the greatest increase in expression of thymidine kinase in cells exposed to combination cis and radiation, consequently leading to high levels of cleaved PARP and decreased levels of phosphorylated ERK in cells exposed to the combination of cis, radiation, and acyclovir (Figure 2). In contrast, the control cell line, HK-1, did not express the enzymatic variation on western blot analysis seen with C666-1. HK-1 also failed to show a difference in cell death.

**Figure 1:**
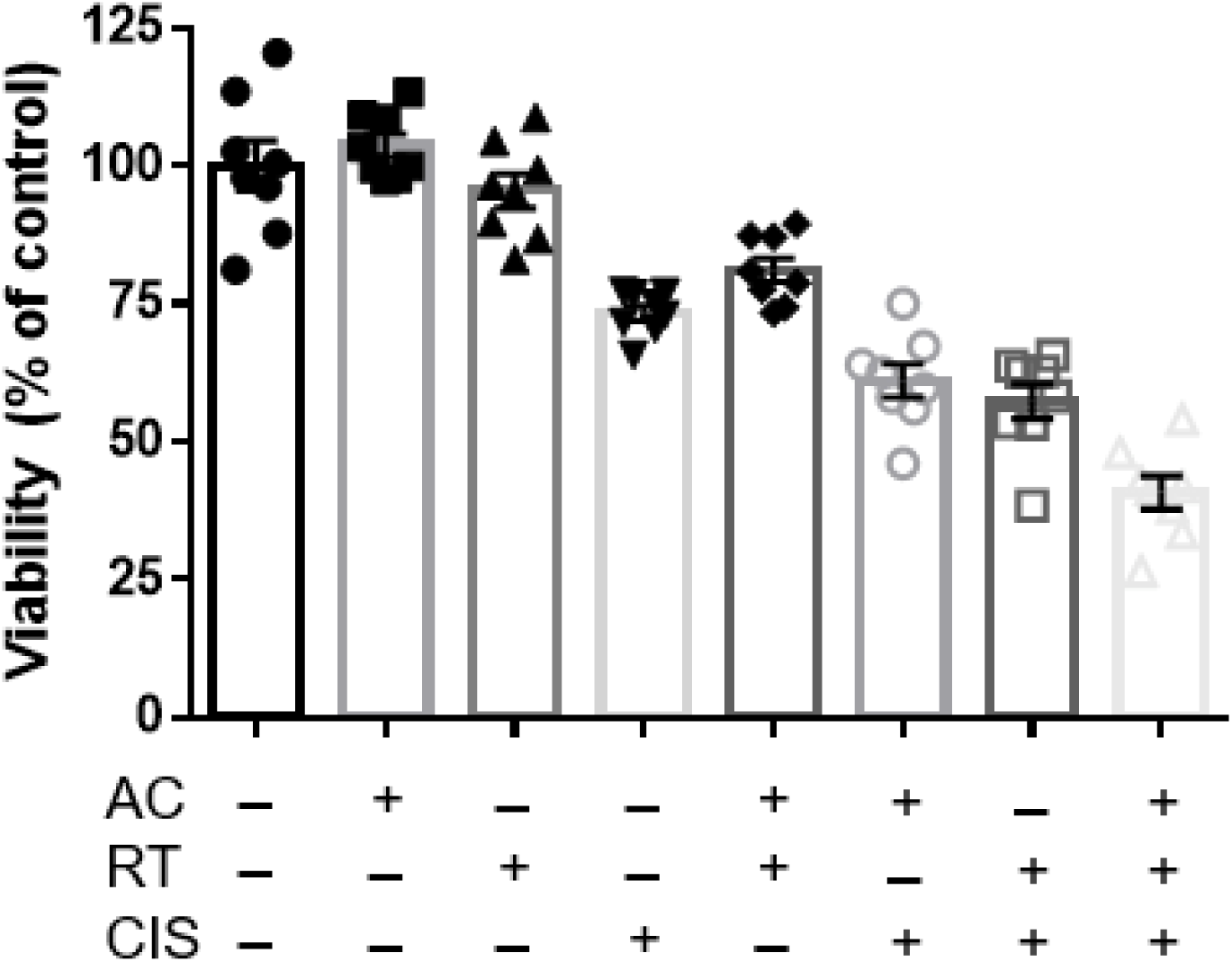
MTT cell viability analysis of EBV-infected C666-1 cells exposed to combinations of no treatment (NT), cisplatin, acyclovir, and 2Gy radiation (RT). The combination of acyclovir and cisplatin reduces cell viability at the 2Gy radiation dosage.

**Figure 2:**
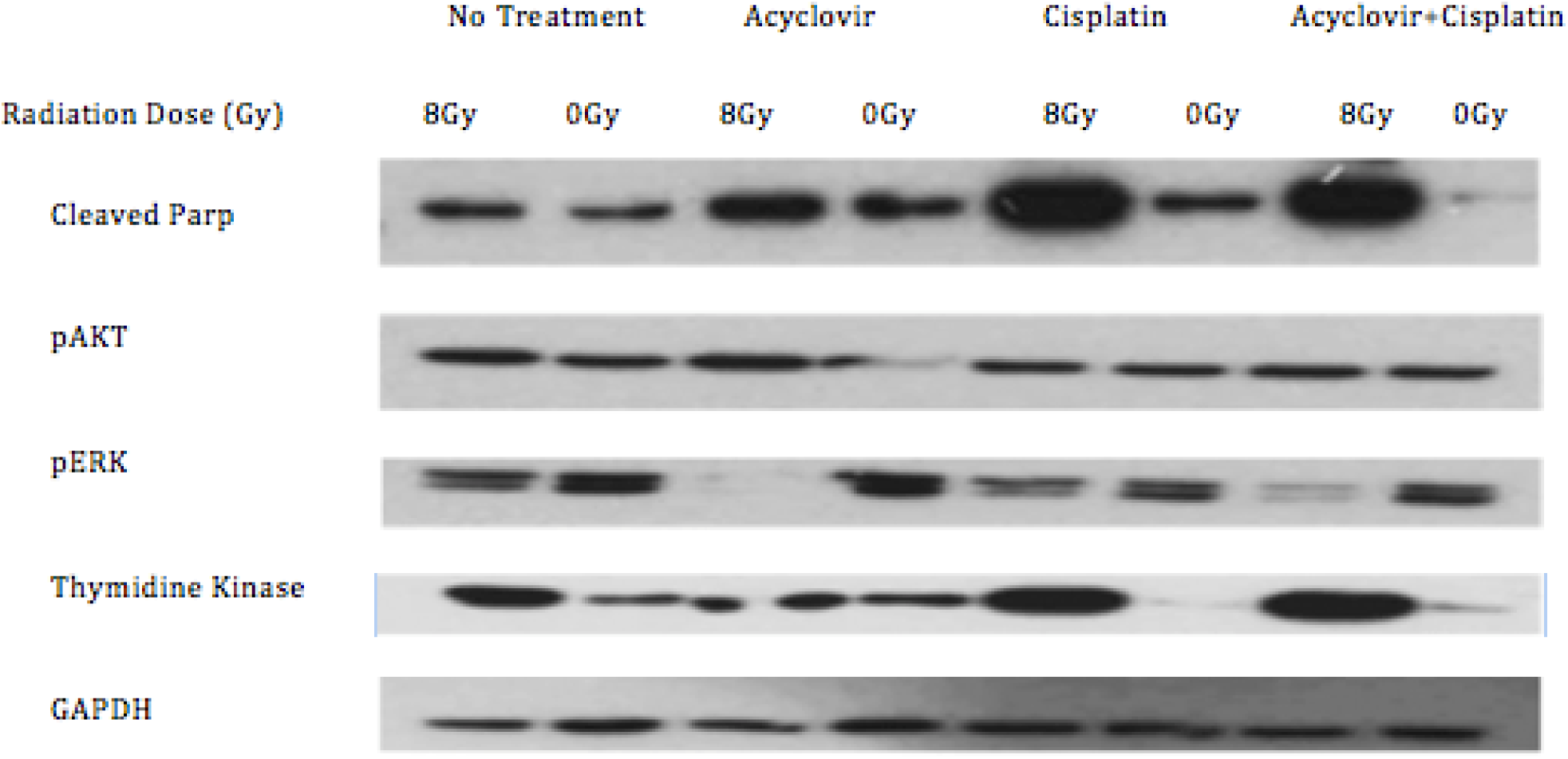
Western blot of the C666-1 cells after receiving no treatment, acyclovir, cisplatin, or radiation. Combination of radiation and cisplatin increases thymidine kinase. Addition of acyclovir increases cParp and decreases pERK.

## Discussion

Recommended treatment for advanced NPC at the time this study was conducted was concurrent chemoradiation with cis followed by adjuvant cis and 5-FU^5^. New data has established induction gemcitabine and cis, followed by cis-RT as another standard of care^9^. This does not alter the relevance of the current study, however, since the backbone is still cis-RT based.

It has been well-established that plasma EBV DNA titers correlate with NPC stage and outcomes. In fact, the HN-001 randomized phase II/III clinical trial assessing risk-adapted treatment of patients with NPX based on EBV DNA titers, is currently in progress (NCT02135042). Targeting virally infected cells has emerged as a novel therapeutic strategy^10^. Although a promising approach, obstacles in efficacy arise as nucleoside-analogues are pro-drugs that can only be activated in the lytic phase by virus specific protein kinases not present in the latent phase^11^. While induction of the lytic phase is possible, data on the efficacy of this technique and synergy with nucleoside analogues is limited in the literature. We strove to establish the critical role of chemoradiation in inducing a lytic switch in latent type II EBV cells, along with the synergestic effect with acyclovir in inducing cell death^10^. Our data shows that the addition of acyclovir to standard of care chemoradiation leads to maximal tumoricidal effects in the C666-1 cells while sparing the control HK-1 line. Increases in TK expression demonstrates a switch into the lytic phase with resultant efficacy of acyclovir as shown by increased levels of cPARP and decreased levels of pERK and subsequent cell death via apoptosis. This synergistic interaction was consistent in enhancing *in vitro* anti-tumor effects across multiple experiments in the C666-1 cells, while sparing the control HK-1 cell line. The synergistic anti-tumor effects of acyclovir, cisplatin, and radiation combination should next be studied with *in vivo* mouse models.

